# Genetic and Environmental Contributions to The Association Between Violent Victimization and Major Depressive Disorder

**DOI:** 10.1101/164194

**Authors:** Nicholas Kavish, Eric J. Connolly, Brian B. Boutwell

## Abstract

Research suggests victims of violent crime are more likely to suffer from major depressive disorder (MDD) compared to non-victims. Less research has utilized longitudinal data to evaluate the directionality of this relationship or examined the genetic and environmental contributions to this association across the life course. The current study evaluated 473 full-sibling pairs and 209 half-sibling pairs (*N* = 1,364) from the National Longitudinal Survey of Youth (M_age_ = 20.14, SD = 3.94). Cross-lagged models were used to examine the directionality of effects between violent victimization and MDD over time. Biometric liability models were used to examine genetic and environmental influences on single and chronic violent victimization and MDD. Violent victimization was associated with increases in MDD during late adolescence, but MDD was more associated with increased risk for violent victimization across young adulthood. Biometric analysis indicated that 20% and 30% of the association between MDD and single and chronic victimization, respectively, was accounted for by common genetic influences. Results from the current study suggest individuals who exhibit symptoms of MDD are at higher risk for chronic victimization rather than developing MDD as a result of victimization. Shared genetic liability accounted for between 20 to 30% of this longitudinal relationship.

## Genetic and environmental contributions to the association between violent victimization and Major Depressive Disorder

In 2015, it was estimated that over 2.7 million U.S. residents age 12 and older experienced a violent victimization (Truman & Morgan, 2016). Based on annual health care expenditures and lost productivity estimates, the economic toll of violence against women alone (i.e., rape, physical assault, and murder) is more than 8 billion dollars annually (Max, Rice, Finkelstein, Bardwell, & Leadbetter, 2004). It is clear that violent victimization is a serious public health concern. At the same time, national reports show that depressive disorders are among the most common mental health illnesses in the U.S. and affect close to 40 million U.S. adults age 18 and older every year (Kessler, Chiu, Demler, & Walters, 2005). Research suggests that the yearly economic burden of individuals with major depressive disorder (MDD) is between 173 and 210.5 billion dollars (in 2005; Greenberg, Fournier, Sisitsky, Pike, & Kessler, 2015). Taken together, there is little doubt that violent victimization and MDD take a large toll on individual victims, their families, and society at large. However, despite theoretical and empirical associations between them (e.g., Barchia & Bussey, 2010; Hawker & Boulton, 2000), research examining the directionality of the association and underlying mechanisms that explain the link between violent victimization and MDD across the life course is scarce.

A substantial body of literature has documented a positive association between victimization and MDD (e.g. Barchia & Bussey, 2010; Hawker & Boulton, 2000; Kaltiala-Heino, Rimpelä, Marttunen, Rimpelä, & Rantanen, 1999; Ranta, Kaltiala-Heino, Pelkonen, & Marttunen, 2009; Slee, 1995). Specifically, violent victimization has been associated with increased symptoms of depression (Boney-McCoy & Finkelhor, 1995; Kilpatrick et al., 2003). For example, a nationally representative sample of adolescents (ages 12-17) reported that exposure to violence increased the risk of a major depressive episode (Kilpatrick et al., 2003).

Similarly, a cross-sectional study of over one thousand adult, family practice patients found that being a victim of interpersonal violence was linked to endorsing more depressive symptoms (Porcerelli et al., 2003). Based on this growing body of research, the standard assumption regarding the relationship between victimization and MDD is that there is a state dependent relationship. That is, individuals who are unfortunate enough to be violently victimized are thought to go on to develop MDD as a direct result of their victimization (Barchia & Bussey, 2010; Boney-McCoy & Finkelhor, 1995; Hawker & Boulton, 2000; Kilpatrick et al., 2003).

Indeed, it seems reasonable to suggest that being victimized, especially violently, could result in the emergence of MDD in many individuals. However, other studies show that not everyone develops MDD following a violent victimization (e.g., DeMaris & Kaukinen, 2005; Vaske, Makarios, Boisvert, Beaver, & Wright, 2009).

A second possible hypothesis for the link between violent victimization and MDD is the population heterogeneity hypothesis. A population heterogeneity perspective suggests that some individuals have characteristics that make them more susceptible to victimization than others, and make some individuals more, or less, likely to develop MDD than others. An important point to consider when evaluating individual differences between violent victimization and MDD concerns the possibility of genetic influences (Barnes et al., 2014). It is now well established that variation across virtually all human phenotypes is at least partly the result of genetic differences in the population (Polderman et al., 2015). Neither violent victimization nor MDD are exempted from this pattern of results, as bodies of research have found that both phenotypes are partly heritable. The moderate heritability of depression has been firmly established by a long line of research (Johnson, McGue, Gaist, Vaupel, & Christensen, 2002; Kendler, Gatz, Gardner, & Pedersen, 2006; Sullivan, Neale, & Kendler, 2000). Although somewhat more so in men, depression is heritable in both sexes and is heritable across the lifespan (Kendler, Gatz, Gardner, & Pedersen, 2006). Furthermore, depression has been found to be heritable in both community and clinical populations (Sullivan, Neale, & Kendler, 2000). These findings suggest that genetic differences may help explain some of the individual variation in the mechanisms, such as neurotransmitter levels and the sensitivity of receptor cells for those neurotransmitters to traumatic events, within the broader population.

More recently, an emerging line of research has also found that being victimized is partly heritable (Beaver, Boutwell, Barnes, & Cooper, 2009; Boutwell et al., 2017; Shakoor et al., 2015). DiLalla and John (2014) found a moderate heritability for “receiving aggression” in 5-year-olds. Shakoor and colleagues (2015) report a modest heritability estimate for bullying victimization in late childhood, and importantly for the current study, report a shared genetic risk for both victimization and paranoid symptoms. Violent victimization has also been found to be heritable in adults, with an even higher heritability estimate for chronic victimization (Beaver, Boutwell, Barnes, & Cooper, 2009). It is important to note that, when describing victimization as heritable, we are not suggesting that there are genes directly causing other people to victimize an individual. Rather, this line of research suggests that there are genetic effects on personal characteristics (e.g., behaviors or personality traits) that elevate an individual’s risk for being victimized.

Given the evidence pertaining to the heritability of victimization and MDD, it is possible that some of the phenotypic covariance between both phenotypes may be accounted for by common additive genetic influences (Barnes, Boutwell, Beaver, Gibson, & Wright, 2014). Put another way, a genetic correlation may exist between violent victimization and MDD over time that explains individual differences in exposure and response to victimization. At least some support for this hypothesis has been found in prior research showing evidence of a shared genetic vulnerability for victimization and other psychopathology including paranoid symptoms (Shakoor et al., 2015), anxiety (Brendgen, Girard, Vitaro, Dionne, & Boivin, 2015), and aggression (Brendgen et al., 2011). However, despite a rapidly accumulating body of evidence in this area of developmental psychopathology, less attention has been given to examining the longitudinal relationships between violent criminal victimization and MDD from adolescence to emerging adulthood, when risk for criminal victimization (Truman & Planty, 2012) and the emergence of MDD symptomology in males and females (Hankin et al., 1998) increases. As a result, the current study seeks to address this gap in the literature by examining the longitudinal relationships between violent victimization and MDD over time in a population-based sample of sibling pairs.

## CURRENT STUDY

The current study analyzes longitudinal sibling data to examine the reciprocity of the association between violent victimization and MDD across time-points. It also evaluates differences in additive genetic and environmental influences on the association between single and chronic victimization with MDD over time. Based on previous research, we hypothesize that violent victimization will be more strongly associated with increases in risk for MDD than MDD with increases in risk for violent victimization. Support for this hypothesis would align with the state dependence hypothesis of victimization, which holds that psychopathology, such as MDD symptomology, develops primarily because of traumatic social life experiences. Evidence in opposition to this hypothesis whereby MDD is more strongly associated with subsequent violent victimization would offer support for the population heterogeneity hypothesis of victimization, which argues that there may be relatively stable dispositional characteristics found in some individuals with chronic depressive symptoms, which make these individuals more vulnerable or places them at risk of experiencing a violent victimization. With respect to the magnitude of common additive genetic and environmental influences on violent victimization and MDD, we hypothesize that variation in liability for chronic victimization will account for more genetic variance in MDD than single victimization owing to selective gene-environment processes.

## METHODS

### Sample

Data analyzed in the current study were drawn from the Child and Young Adult sample of the National Longitudinal Survey of Youth (CNLSY). The CNLSY consists of a sample of youth born to a nationally representative sample of females from the National Longitudinal Survey of Youth 1979 (NLSY79). All CNLSY children born to women from the NLSY79 over their life have been assessed biennially (Chase-Lansdale, Mott, Brooks-Gunn, & Phillips, 1991). When children have reached the age of 15, they were administered the Youth Adult (YA) self-report survey at each survey wave. The YA survey allows children to respond to several different age-appropriate sensitive questions in a private self-reporting format. Thus, respondents are asked to report on topics including, delinquent behavior, family relationships, personal attitudes, and violent victimizations. Retention rates for children from the CNLSY from 1986 to 2012 have been above 70%.

Since the 2012 survey wave, nearly 5,000 females from the NLSY79 have given birth to over 11,500 children. Kinship links have been established by Rodgers and colleagues (2016) using explicit indicators of sibling status (e.g., twin sibling, full-sibling, half-sibling, or adopted sibling) based on questionnaires first administered during the 2006 survey wave. Since 2006, 16,083 sibling pairs (100% of possible pairs) from the CNLSY have been identified using the kinship links (Hadd & Rodgers, 2017). Based on the richness of these data, several researchers have used the kinship links (indicators of additive genetic relatedness between siblings) to conduct behavioral genetic and sibling comparison analyses on a wide range of topics including antisocial behavior (Van Hulle et al., 2009), delinquency (Connolly & Beaver, 2014; Harden, Quinn, & Tucker-Drob, 2012), intelligence (Rodgers, Rowe, May, 1994), maternal smoking (D’Onofrio et al., 2008), maternal nutrition (Connolly & Beaver, 2015), and violent victimization (Boutwell et al., 2017) (see Rodgers et al., 2016, for background and summary). Researchers interested in taking advantage of the CNLSY kinship pairs can find documentation and code for the links at http://liveoak.github.io/NlsyLinks/.

The current study takes advantage of the kinship links and uses data on respondents who had at least one full- or half-sibling with complete data on violent victimization and MDD. Only one sibling pair per household, randomly selected, was included in the sample. The analytic sample was restricted to full- and half-sibling pairs because there were too few identical twin pairs (*n* = 7) to include in the analysis. The final sample therefore included *n* = 473 full-sibling pairs and *n* = 209 half-sibling pairs (*N* = 1,364 siblings).

### Measures

#### Major Depressive Disorder

MDD was assessed by the seven-item Center for Epidemiologic Studies Depression Scale (CESD-D) short form (CES-D-SF). Respondents were asked during the 2004, 2006, 2008, 2010, and 2012 survey wave to report how often in the past week they had suffered from each of the following symptoms: (1) poor appetite, (2) trouble focusing, (3) feeling depressed, (4) everything taking extra effort, (5) restless sleep, (6) saddened, and (7) unable to get “going”. Response categories ranged from 0 (*rarely, none of the time, 1 day*) to 4 (*most, all the time, 5-7 days*). Responses were summed to create scales of depression at each survey wave. Scales demonstrated adequate internal consistency across time with Cronbach’s alphas ranging from .68 to .71 and all items loading highly on a single common factor at each survey wave. Previous research analyzing data from the NLSY79 has shown that the CES-D-SF demonstrates better psychometric properties compared to the CES-D with higher internal consistency and better unidimensionality (Levine, 2013). Moreover, evidence from prior psychometric research indicates that a CES-D-SF cut-off score of 8 or more has acceptable specificity with the standard CES-D cutoff score of 16 or more (Levine, 2013), which would satisfy the DSM-V criteria for MDD (American Psychiatric Association, 2013). As such, to assess risk for MDD at each survey wave, CESD-D-SF scales were dichotomized such that 0 = score of 7 or less and 1 = score of 8 or more. A cumulative scale of MDD across survey was also created by adding together all dichotomized measures to examine the frequency of MDD over time. Table 1 presents the percentage of respondents classified as suffering from MDD at each survey wave.

**Table 1.** Descriptive Statistics

|  | Mean/Percent | SD | Min | Max | n | Skewness | Kurtosis |
| --- | --- | --- | --- | --- | --- | --- | --- |
| Major Depression<sub>2004</sub> | 18.97% | - | 0 | 1 | 1532 | .86 | 2.40 |
| Major Depression<sub>2006</sub> | 19.60% | - | 0 | 1 | 1910 | .91 | 3.69 |
| Major Depression<sub>2008</sub> | 17.95% | - | 0 | 1 | 2026 | .77 | 2.19 |
| Major Depression<sub>2010</sub> | 15.64% | - | 0 | 1 | 1984 | .99 | 3.00 |
| Major Depression<sub>2012</sub> | 14.60% | - | 0 | 1 | 1080 | 1.05 | 3.57 |
| Major Depression<sub>2004-2012</sub> | .85 | 1.20 | 0 | 5 | 1330 |  |  |
| Violent Victimization<sub>2004</sub> | 8.56% | - | 0 | 1 | 1106 |  |  |
| Violent Victimization<sub>2006</sub> | 12.50% | - | 0 | 1 | 1850 |  |  |
| Violent Victimization<sub>2008</sub> | 6.50% | - | 0 | 1 | 2042 |  |  |
| Violent Victimization<sub>2010</sub> | 4.00% | - | 0 | 1 | 1600 |  |  |
| Violent Victimization<sub>2012</sub> | 4.17% | - | 0 | 1 | 1048 |  |  |
| Single Violent Victimization | 8.04% | - | 0 | 1 | 1364 |  |  |
| Chronic Violent Victimization | 2.31% | - | 0 | 1 | 1364 |  |  |
| Age | 20.14 | 3.94 | 15 | 30 | 1364 |  |  |
| Black or Hispanic | 46% | - | 0 | 1 | 1364 |  |  |
| Male | 51% | - | 0 | 1 | 1364 |  |  |

*Note:* Skewness and Kurtosis are for the CESD (depression inventory) prior to dichotomization.

#### Violent Victimization

Violent victimization was first assessed during the 2004 survey wave and then again during the 2006, 2008, 2010, and 2012 survey waves by asking respondents if they had been the victim of a violent crime (e.g., physical or sexual assault, robbery or arson) since the date of their last interview. To reduce social desirability bias and encourage accurate reporting, respondents were asked to report whether they had been the victim of violent crime in the YA survey, which is administered in a private self-reporting format. Response categories were 0 = no and 1 = yes. To assess differences in MDD over time between single and chronic victims, two additional measures were created. The first measure was created to identify respondents who had only experienced one victimization from 2004 to 2012. This measure of single victimization was coded 0 = no victimization and 1 = one victimization. The second measure created was designed to identify respondents who had experienced more than one violent victimizations from 2004 to 2012. The chronic victimization measure was coded 0 = none or 1 victimization and 1 = two or more victimizations. Table 1 reports the percentage of respondents who experienced a single victimization and more than one victimization. Overall, 8.04% of the sample reported one violent victimization, while 2.31% reported two or more violent victimizations.

#### Statistical Controls

Age, race, and sex were controlled for in all analyses. Age was measured by the age of respondents (in years) during the 2004 survey wave. Race was measured by a dichotomous variable where 0 = Non-Black and Non-Hispanic and 1 = Black or Hispanic. Sex was measured by a dichotomous variable where 0 = female and 1 = male. Table 1 shows that the average age of respondents was 20 years old, 46% were Black or Hispanic, and 51% were male.

## PLAN OF ANALYSIS

The plan of analysis was carried out in three interrelated steps. First, an autoregressive cross-lagged model was fit to the data to examine whether victimization at earlier waves predicted increases in probability of MDD at later waves while controlling for stability in both measures and statistical controls. Autoregressive cross-lagged models offer a unique advantage over other longitudinal modeling approaches, such as latent growth curve modeling and group-based trajectory modeling, in that they allow for the estimation of the reciprocal effects on change between two variables over time, while maintaining temporal order (Selig & Little, 2012). While latent growth curve modeling and group-based trajectory modeling provide evidence of correlated within-individual and group change over time, neither approach can provide estimates of the bidirectional associations (i.e., whether victimization is more strongly associated with risk for future MDD than MDD is associated with risk for future victimization). All autoregressive cross-lagged models were estimated in M*plus* 7.4 (Muthén & Muthén, 2012) using a weighted least squares variance estimator (WLSMV) which is appropriate for latent variables with binary or ordinal properties. Model fit was assessed using the comparative fit index (CFI), Tucker-Lewis index (TLI), and root mean square error of approximation (RMSEA). As recommended (Hu & Bentler, 1999), the following cut-off points were used to assess acceptable model fit: CFI ≥ .90, TLI ≥ .90, and RMSEA ≤ .08. Standard errors were also adjusted for non-independence since several respondents coming from the same household were nested within the analytic sample.

Second, between-sibling correlations were calculated to examine the concordance between siblings for MDD and violent victimization. Between-sibling correlations are traditionally first calculated before biometric model fitting approaches are used to investigate whether there is evidence of genetic influence on a measure. Evidence for genetic influence would appear if siblings who share more genetic material have stronger between-sibling resemblance or concordance on a measure compared to siblings who share less genetic material. With regards to the present study, if full-sibling pairs (who share, on average, 50% of their genetic material with one another) demonstrate stronger concordance for MDD and violent victimization compared to half-siblings (who share, 25% of their genetic material with one another) then this can be interpreted as evidence of genetics partly explaining variation in risk for MDD and violent victimization. Since the cumulative measure of MDD analyzed in this step of the analysis was a categorical variable, and both measures of single and chronic victimization were binary variables, different types of correlations were calculated to assess between-sibling concordance for all three variables. Spearman’s correlation coefficients were calculated for the cumulative measure of MDD and used to assess differences in concordance between full- and half-sibling pairs, while tetrachoric correlation coefficients were calculated for single and chronic victimization.

Third, univariate liability-threshold models were estimated to partition the total observed variance in liability for MDD and violent victimization into unstandardized (which were subsequently converted into standardized) estimates of variance attributable to additive genetic (symbolized as A), shared environmental (symbolized as C), and non-shared environmental (symbolized as E) influences (Neale & Cardon, 1992; Prescott, 2004). The magnitude of A represents the amount of variance in liability accounted for by additive genetic influences, while the magnitude of C represents the amount of variance in liability accounted for by shared environmental experiences that make siblings like one another for MDD and risk for violent victimization. The magnitude of E represents the amount of variance in liability accounted for by environmental experiences that are not shared between siblings, but are unique to each sibling and create differences in risk for MDD and violent victimization between them. Measurement error is also included in the E estimate (Plomin, DeFries, Knopik, & Neiderhiser, 2013). After evaluating parameter estimates from the best fitting univariate ACE models, bivariate liability-threshold models were estimated to determine if genetic and environmental influences on MDD were shared with single and chronic victimization (see Figure 1). The variance in liability for MDD was therefore decomposed into additive genetic and environmental components common to single and chronic victimization and unique of single and chronic victimization. The proportion of total variance in liability for MDD accounted for by genetic influences shared with single and chronic victimization was calculated using the following algebraic equation:

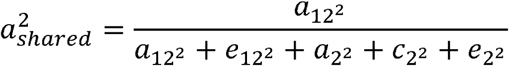

**Figure 1.**
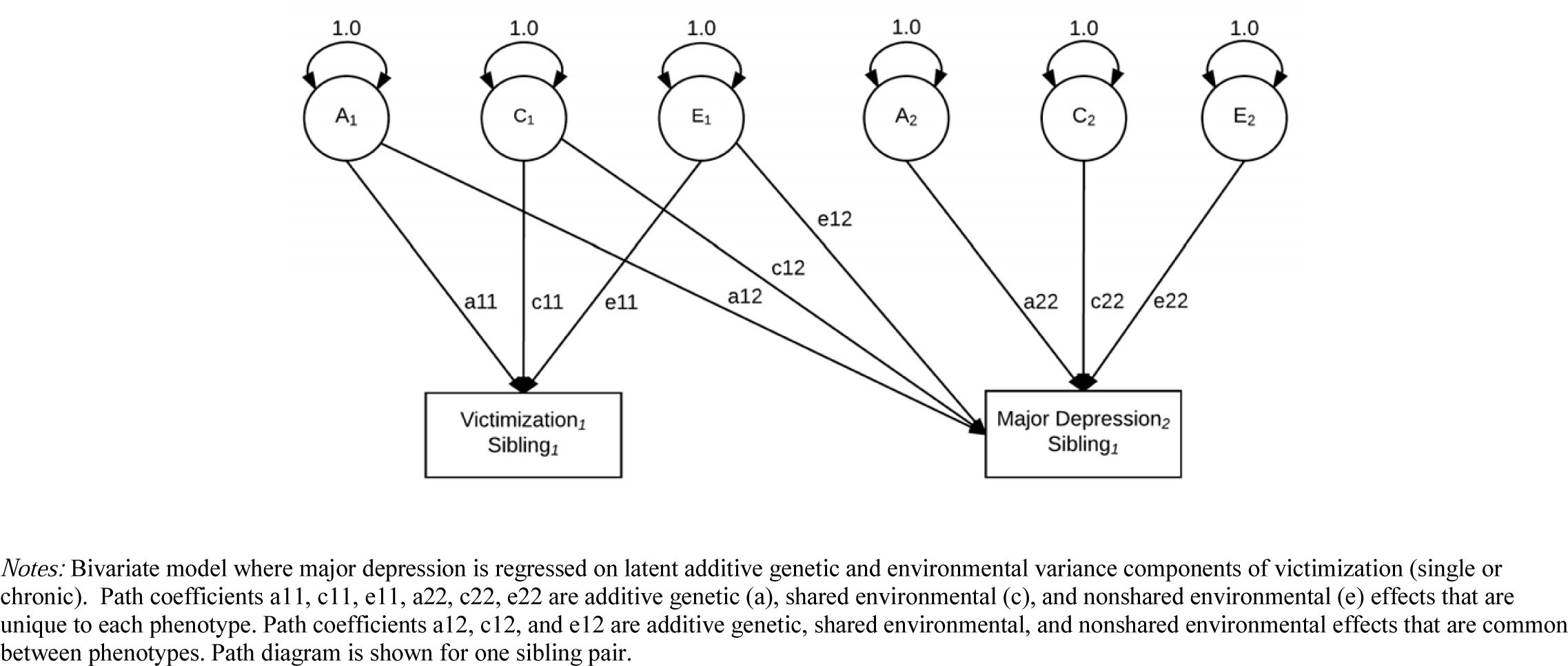
Path Diagram for Bivariate Model.

Model fit for each univariate and bivariate liability-threshold model was evaluated using the following model fit indices: the Satorra-Bentler scaled-difference chi-square test (Δχ^2^) where a nonsignificant change in chi-square indicates that the model with fewer estimated parameters fits the data equally well and is preferred for the sake of parsimony (Satorra & Bentler, 2001); the comparative fit index (CFI) where CFI ≥ .90 indicates an adequate fit; the root mean square error of approximation (RMSEA) where RMSEA ≤ .08 indicates an adequate fit (Hu & Bentler, 1999).

Because we restricted our analysis to only one full- or half-sibling pair per household with complete data on MDD and victimization from 2004 to 2012, the sibling pairs analyzed in the current study are a subset of the 16,083 sibling pairs nested within the CNLSY. Preliminary analyses were conducted to examine whether this resulted in a biased sample. Equal proportions of male and female siblings were included in the analysis, but a higher percentage of Black (30% vs. 27%) and Hispanic (23% vs. 17%) siblings were included in the analysis compared to excluded siblings. There was no substantive difference in age between siblings included and excluded from the current analysis (20.14 vs. 20.87).

## RESULTS

### Phenotypic Analysis

Before estimating a cross-lagged model, phenotypic associations between MDD and violent victimization at each survey wave were examined. Since all measures were binary, tetrachoric correlations were used to examine associations between all variables. Table 2 summarizes the phenotypic associations between MDD and victimization from 2004 to 2012. As can be seen, there was moderate stability in risk for MDD (*rho* = .46 to *rho* = .50, *p* < .001) and violent victimization (*rho* = .28 to *rho* = .51, *p* < .001) across waves. MDD was positively and significantly associated with violent victimization at each survey wave (*rho* = .16 to *rho* = .28, *p* < .001).

**Table 2.** Polychoric and Tetrachoric Phenotypic Correlations for Major Depression and Violent Victimization

|  | 1. | 2. | 3. | 4. | 5. | 6. | 7. | 8. | 9. | 10. |
| --- | --- | --- | --- | --- | --- | --- | --- | --- | --- | --- |
| 1. Major Depression<sub>2004</sub> | - |  |  |  |  |  |  |  |  |  |
| 2. Major Depression<sub>2006</sub> | .48** | - |  |  |  |  |  |  |  |  |
| 3. Major Depression<sub>2008</sub> | .45** | .48** | - |  |  |  |  |  |  |  |
| 4. Major Depression<sub>2010</sub> | .43** | .47** | .50** | - |  |  |  |  |  |  |
| 5. Major Depression<sub>2012</sub> | .39** | .40** | .47** | .46** | - |  |  |  |  |  |
| 6. Violent Victimization<sub>2004</sub> | .22** | .14** | .13** | .09 | .25** | - |  |  |  |  |
| 7. Violent Victimization<sub>2006</sub> | .20** | .21** | .27** | .16** | .21** | .51** | - |  |  |  |
| 8. Violent Victimization<sub>2008</sub> | .10* | .17** | .16** | .21** | .22** | .29** | .28** | - |  |  |
| 9. Violent Victimization<sub>2010</sub> | .09 | .15** | .20** | .26** | .17* | .31** | .24** | .37** | - |  |
| 10. Violent Victimization<sub>2012</sub> | .09 | .19** | .21** | .19** | .28** | .24* | .30** | .30** | .42** | - |

Notes: \*\* $p < .001$ ; \* $p < .01$

### Cross-Lagged Analysis

After establishing that MDD and violent victimization were associated with one another from 2004 to 2012, we estimated a bidirectional cross-lagged model to control for stability in both measures and assess the relationship between variables over time. As shown in Table 3, tests for factorial invariance of MDD over time revealed that strong invariance (equality of factor loadings and measurement error) did not greatly reduce model fit, thus suggesting that the same construct was likely being measured across assessment waves. The final specified model included controls for age, race, and sex. Overall, the final model provided a close fit to the data (χ^2^ = 191.87, *p* < .001, CFI = .95, TLI = .91, RMSEA = .04 [95% CI: .02-.06]). Standardized coefficients from the specified model are presented in Figure 2. As shown, standardized path coefficients revealed that while violent victimization in 2004 was associated with increased risk for MDD in 2006 (*β* = .09, *p* < .01), MDD was more strongly, and consistently, associated with subsequent increases in risk for violent victimization over time. Specifically, MDD was associated with increases in victimization from Wave 1 to Wave 2 (*β* = .13, *p* < .001), Wave 2 to Wave 3 (*β* = .15, *p* < .001), Wave 3 to Wave 4 (*β* = .18, *p* < .001), and Wave 4 to Wave 5 (*β* = .16, *p* < .001).

**Table 3.** Model Fit Indices for Configural, Weak, and Strong Factorial Invariance

| Model | $\chi^2$ | $df$ | CFI | RMSEA (95% CI) | $\Delta\chi^2$ | $\Delta df$ | $p$ |
| --- | --- | --- | --- | --- | --- | --- | --- |
| Configural | 159.41 | 48 | .96 | .05 | - | - | - |
| Weak | 188.19 | 53 | .95 | .04 | 14.32 | 5 | .03 |
| Strong | 191.57 | 58 | .95 | .04 | 6.79 | 5 | .19 |

**Figure 2.**
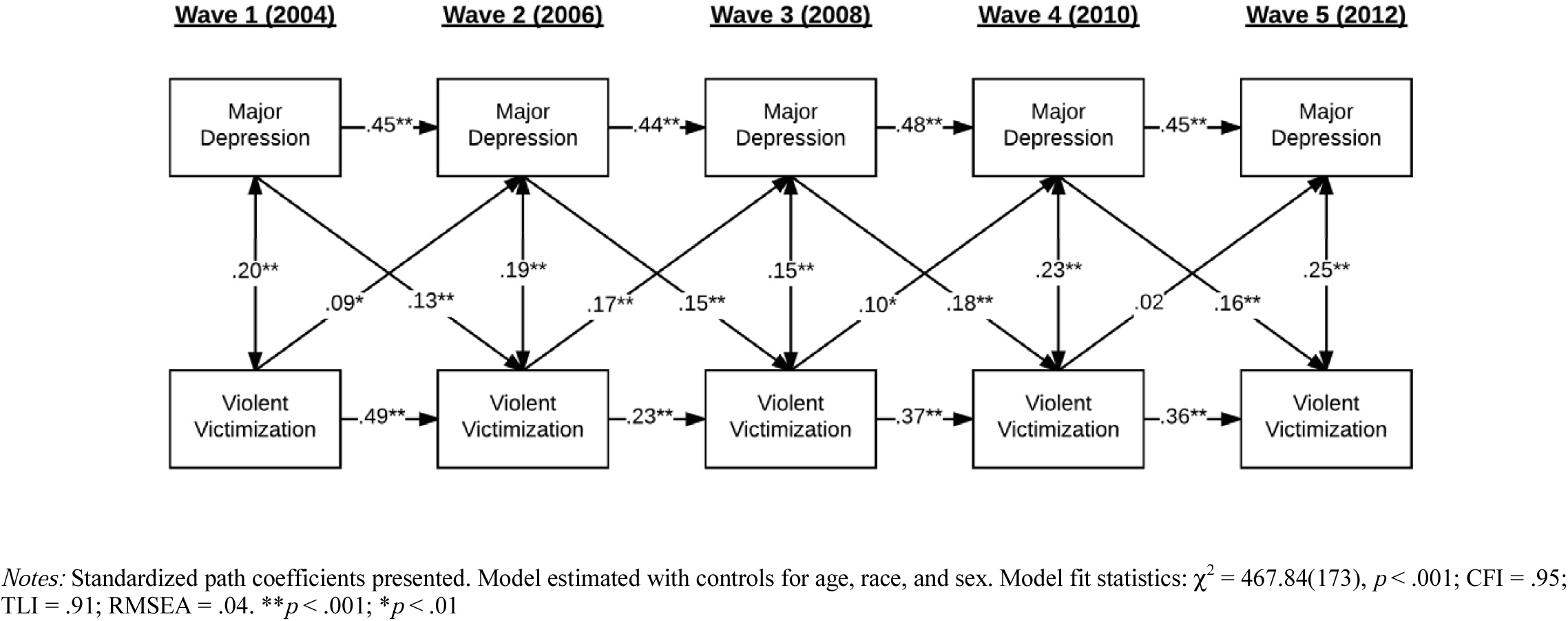
Cross-Lagged Model for Major Depression and Violent Victimization.

### Behavioral Genetic Analysis

To better understand the underlying mechanisms involved in the association between MDD and victimization over time, behavioral genetic analyses were conducted. Table 4 summarizes the within sibling pair correlations for the cumulative measure of MDD, single violent victimization, and chronic violent victimization. Full-siblings demonstrated stronger concordance for major depressive episodes (*r* = .31, *p* < .01), single victimization (*rho* = .34, *p* < .01), and chronic victimization (*rho* = .43, *p* < .01) compared to half-siblings. Within sibling pair correlations for full-siblings were not more than double the size of correlation for half-siblings, thus suggesting that dominance genetic effects (D) did not warrant further investigation. The results thus suggested that additive genetic influences partly accounted for variance in liability for MDD and violent victimization.

**Table 4.**
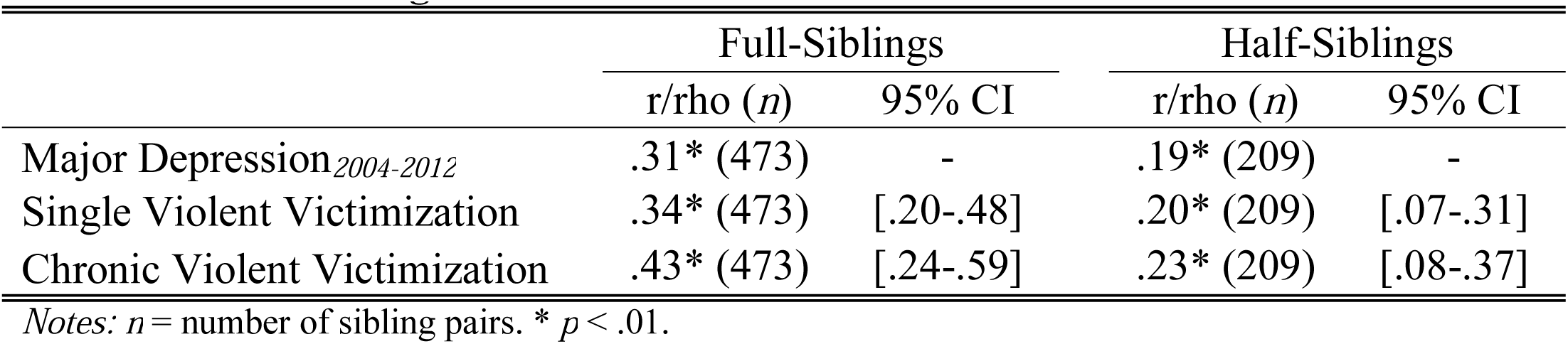
Cross-Sibling Correlations

|  | Full-Siblings |  | Half-Siblings |  |
| --- | --- | --- | --- | --- |
|  | r/rho (*n*) | 95% CI | r/rho (*n*) | 95% CI |
| Major Depression<sub>2004-2012</sub> | .31* (473) | - | .19* (209) | - |
| Single Violent Victimization | .34* (473) | [.20-.48] | .20* (209) | [.07-.31] |
| Chronic Violent Victimization | .43* (473) | [.24-.59] | .23* (209) | [.08-.37] |

Notes: *n* = number of sibling pairs. \* $p < .01$ .

Univariate ACE models were then estimated to identify the magnitude of additive genetic and environmental influences on all MDD and victimization. All estimated univariate models and standardized parameter estimates are summarized in Table 5, with the best-fitting model bolded. As shown, the best-fitting model for MDD was an ACE model (CFI = .89, RMSEA =.06) where 38% of the variance in liability was attributable to additive genetic influences, 16% of the variance in liability was attributable to shared environmental influences, and 46% of the variance in liability was attributable to nonshared environmental influences. The best-fitting model for single violent victimization was an AE model (Δχ^2^ = 1.18, *p* = .41, CFI = .92, RMSEA =.06) where 36% of the variance in liability for experiencing a single victimization was attributable to additive genetic influences, while 64% of the variance in liability was attributable to nonshared environmental influences. An AE model also fit the data more closely than a full ACE model for chronic victimization (Δχ^2^ = .07, *p* = .26, CFI = .88, RMSEA =.06).

**Table 5.** Parameter Estimates from Univariate ACE Models

| | A | C | E | $\Delta\chi^2$ | $\Delta df$ | $p$ | CFI | RMSEA |
| --- | --- | --- | --- | --- | --- | --- | --- | --- |
| Major Depression |  |  |  |  |  |  |  |  |
| ACE | **.38\*\***<br>[.28-.51] | **.16\***<br>[.04-.26] | **.46\*\***<br>[.39-.62] | - | - | - | **.89** | **.06** |
| AE | .49**<br>[.29-.62] | .00<br>[.00-.00] | .51**<br>[.38-.71] | 25.71* | 4 | <.01 | .80 | .07 |
| CE | .00<br>[.00-.00] | .24**<br>[.14-.39] | .76**<br>[.61-.86] | 49.03* | 4 | <.01 | .77 | .08 |
| E | .00<br>[.00-.00] | .00<br>[.00-.00] | 1.00***<br>[1.00-1.00] | 104.55** | 5 | <.001 | .60 | .09 |
| Single Violent Victimization |  |  |  |  |  |  |  |  |
| ACE | .32**<br>[.19-.44] | .08<br>[-.01-.11] | .60**<br>[.51-.86] | - | - | - | .90 | .07 |
| AE | **.36\*\***<br>[.25-.50] | **.00**<br>[.00-.00] | **.64\*\***<br>[.50-.75] | **1.18** | **1** | **.41** | **.92** | **.06** |
| CE | .00<br>[.00-.00] | .10*<br>[.01-.18] | .90**<br>[.82-.99] | 37.81** | 1 | <.001 | .74 | .08 |
| E | .00<br>[.00-.00] | .00<br>[.00-.00] | 1.00***<br>[1.00-1.00] | 92.61** | 2 | <.001 | .69 | .09 |
| Chronic Violent Victimization |  |  |  |  |  |  |  |  |
| ACE | .49**<br>[.36-.69] | .01<br>[-.02-.04] | .50**<br>[.32-.62] | - | - | - | .86 | .07 |
| AE | **.51\*\***<br>[.38-.74] | **.00**<br>[.00-.00] | **.49\*\***<br>[.26-.62] | **.07** | **1** | **.26** | **.88** | **.06** |
| CE | .00<br>[.00-.00] | .09*<br>[.01-.13] | .91**<br>[.87-.99] | 47.40** | 1 | <.001 | .65 | .08 |
| E | .00<br>[.00-.00] | .00<br>[.00-.00] | 1.00***<br>[1.00-1.00] | 110.61** | 2 | <.001 | .60 | .10 |

Notes: Standardized ACE parameter estimates presented. 95% confidence intervals presented in brackets. CFI = comparative fit index, RMSEA = root mean square error of approximation. \*\* $p < .001$ ; \* $p < .01$

Standardized parameter estimates from the best-fitting AE model revealed that 51% of the variance in liability for chronic violent victimization over time was attributable to additive genetic influences, while 49% of the variance in liability was attributable to nonshared environmental influences.

Bivariate liability-threshold models were then used to examine genetic and environmental influences on MDD shared with single and chronic victimization. Unstandardized parameter estimates from the best-fitting bivariate models are summarized in Table 6. As can be seen, after calculating the proportion of covariance between MDD and single victimization using the unstandardized parameter estimates, the results revealed that 20% of the variance in liability for MDD was shared with single victimization. The same procedure for MDD and chronic victimization revealed that 30% of the variance in liability for MDD was shared with chronic victimization. Additional standardized estimates for additive genetic and environmental influences of MDD common to, and unique of, single and chronic violent victimization are presented in Figures 3 and 4. Based on the presented estimates, there was a small, albeit significant, nonshared environmental overlap between MDD and both single and chronic victimization over time (1%, 95% CI: .01-.03). The results also show that genetic factors unique to MDD (22%) played a more important role in the association between MDD and single victimization, compared to MDD and chronic victimization (10%).

**Table 6.** Parameter Estimates from Bivariate Models

| | A<sub>1</sub> | E<sub>1</sub> | b<sub>A</sub> | b<sub>E</sub> | A<sub>2</sub> | C<sub>2</sub> | E<sub>2</sub> | $a_{shared}^2$ | 95% CI |
| --- | --- | --- | --- | --- | --- | --- | --- | --- | --- |
| Major Depression /<br>Single Violent Victimization | .58 (.05) | .81 (.04) | .45 (.08) | .06 (.05) | .46 (.20) | .40 (.14) | .65 (.05) | .20* | [.07-.32] |
| Major Depression /<br>Chronic Violent Victimization | .70 (.05) | .71 (.03) | .54 (.06) | .11 (.05) | .40 (.20) | .33 (.17) | .64 (.05) | .30** | [.19-.41] |

Notes: $a_{shared}^2$ = proportion of covariance between major depression and victimization explained by shared additive genetic influences. CI = confidence intervals. \*\* $p < .001$ ; \* $p < .01$

**Figure 3.**
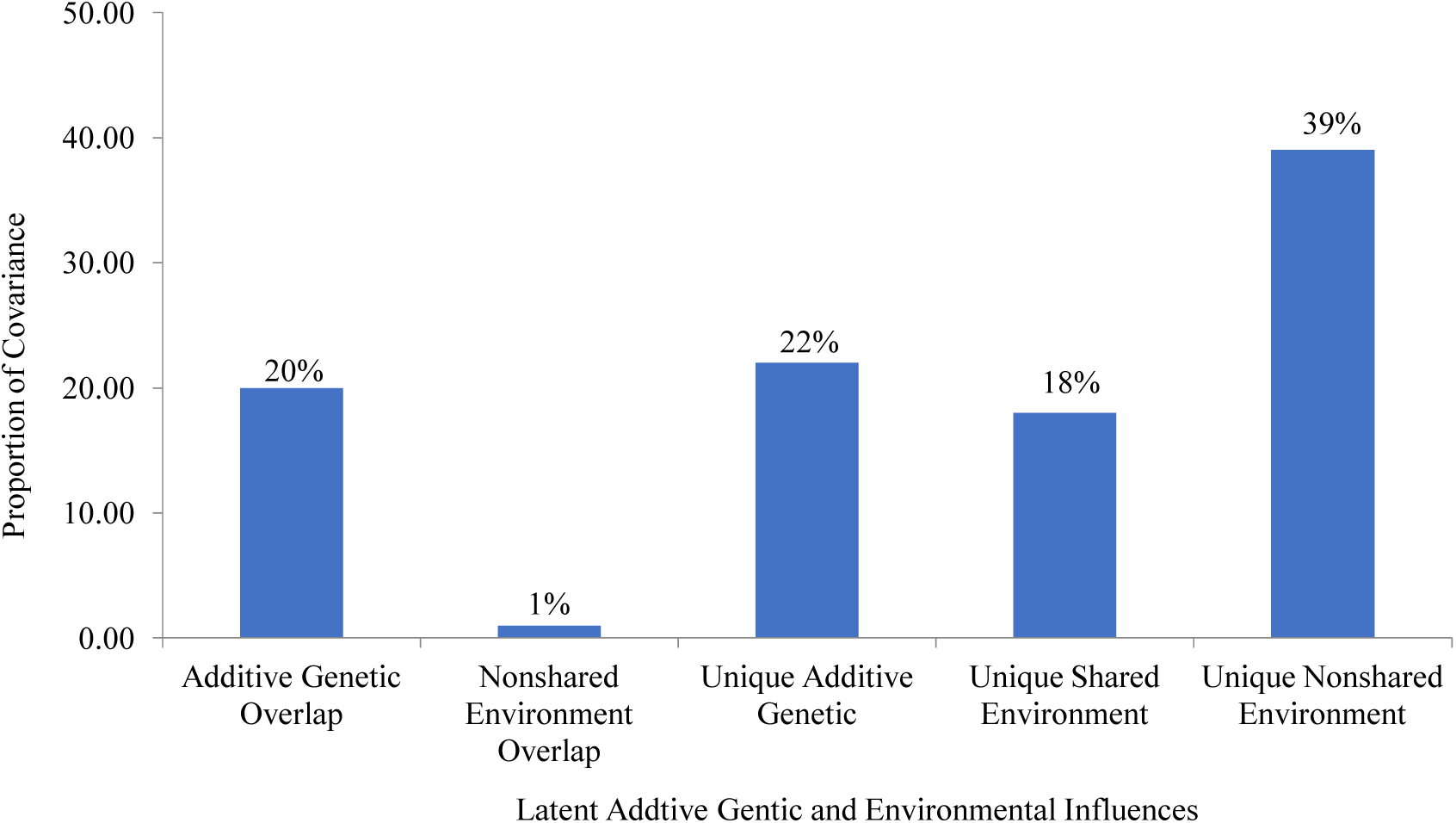
Latent Genetic and Environmental Overlap between Major Depression and Single Violent Victimization.

**Figure 4.**
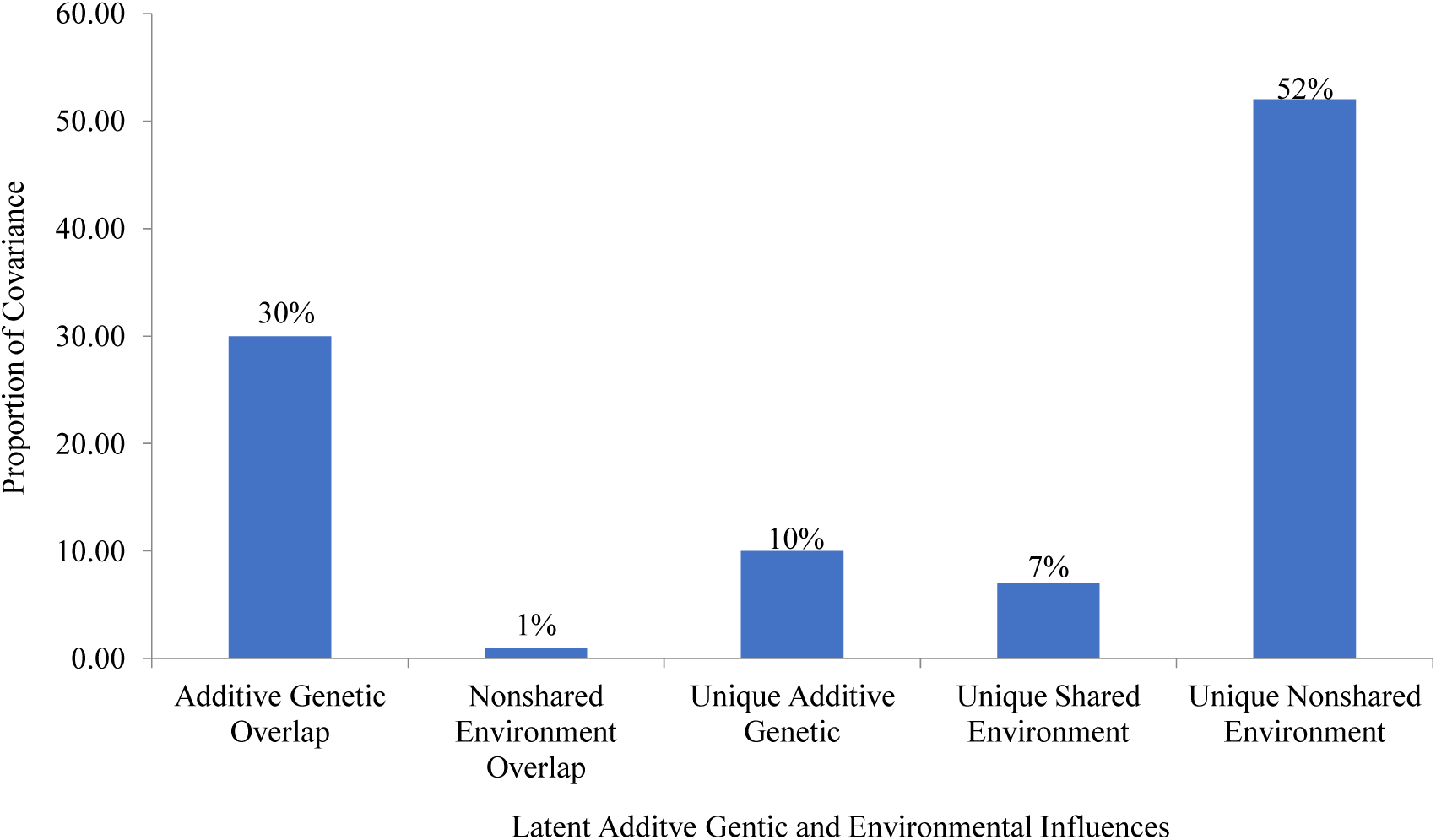
Latent Genetic and Environmental Overlap between Major Depression and Chronic Violent Victimization.

## DISCUSSION

The current study sought to further elucidate the strength and bi-directional relationship between MDD and violent victimization over time. In particular, we hypothesized that violent victimization at a given time point would be more strongly associated with an increased risk for future MDD symptomology. Contrary to this hypothesis, bi-directional cross-lagged analyses revealed that while victimization reported during 2004 was positively associated with MDD during 2006 during late adolescence, MDD was more strongly associated with a higher risk for violent victimization from late adolescence to young adulthood. These results offer preliminary support for a self-selection, or population heterogeneity hypothesis, which suggests that individual traits and behaviors may increase risk some individuals to be exposed to circumstances that elevate their odds of experiencing a violent victimization, compared to others in a population (Beaver et al., 2016). These findings also offer a cautionary note about inferring the directionality of a correlation between two factors measured at a single time point as has been done in much of the prior research on this topic (e.g., Kaltiala-Heino et al., 1999; Kilpatrick et al., 2003).

In the second phase of the analysis, we assessed the magnitude of common genetic and environmental influences on single victimization and MDD as well as chronic victimization and MDD. We hypothesized that there would be a genetic overlap between MDD and violent victimization and that there would be a larger shared genetic component between chronic victimization and MDD than between single victimization and MDD. In line with our hypothesis, bivariate liability models revealed that the genetic covariance in liability between single victimization and frequency of MDD was 20% while the covariation between chronic victimization and frequency of MDD was 30%. Finding for a stronger genetic overlap between chronic victimization and MDD is also consistent with the selection hypothesis of victimization as selection may play a more prominent role in chronic victimization, whereas single victimization has a higher likelihood of happening randomly.

In previous decades, scholars have articulated selection processes by which genetic propensities can contribute to increased risk of exposure to deleterious environments, such as what we observed in the current study (Scarr & McCartney, 1983). In this case, active or evocative gene-environment correlation (*r*GE) could be used to explain why some individuals who suffer from major depressive episodes are more likely to experience chronic violent victimization compared to others (Scarr & McCartney, 1983). To illustrate, active gene-environment correlation occurs when an individual actively selects into an environment based on genetically influenced personality and temperamental traits, while evocative gene-environment correlation takes place when an individual evokes specific behavioral or verbal responses from members of their environment in part because of genetically influenced traits. Based on this logic, individuals with a higher level of genetic liability for developing major depressive disorder may engage in risky behaviors, such as heavy alcohol or illicit drug use, to cope with symptoms which lead them into unsafe environments with antisocial individuals (active *r*GE). Alternatively, individuals with a genetic predisposition for MDD may be victimized by others, including those with a wider range of pathological characteristics besides antisocial features (for example, individuals with Borderline Personality Disorder), because they are perceived as vulnerable targets (evocative *r*GE) due to feelings of worthlessness and excessive guilt. Additional research using other samples are needed to evaluate this possibility and help to further elucidate the specifics of how these selection processes work.

The current study is not without limitations, chief among them being the dichotomous coding of the relevant variables. First, prior researchers in the area of psychopathology have rightly pointed out the potential problems with the use of arbitrary cut-points separating psychopathology from otherwise “normal” variation on some trait (Kotov et al., 2017). Furthermore, in clinical practice, more than a brief inventory of symptoms (i.e., a clinical interview) is generally needed to diagnose MDD. The same limitations apply to our measure of victimization. Though we were able to assess chronic violent victimization over time (broadly defined), this part of our analysis relies on dichotomous indicators of victimization that were asked repeatedly across several waves of data collection. The use of a binary item restricts variation in undesirable ways, and does not allow us to examine victimization across a range of possible experiences (i.e., physical assaults versus sexual assaults). To the extent that our findings replicate with better measures of victimization (and continuously measured psychopathological items) remains an open empirical question.

Pending independent replication, the results of this study suggest a need for more research into the association between violent criminal victimization and psychopathology that can better assess the direction of relationships. Future research will also benefit from examining what specific kinds of psychopathology, as well as which specific symptoms, are most associated with victimization and evaluate how these traits might be increasing an individual’s risk for violent victimization. Additionally, while genetically sensitive designs are not always feasible, the current study re-emphasizes the tenuousness of causally interpreting the associations found between two or more phenotypes without controlling for unobservable genetic confounds (Barnes et al., 2014).

